# Engagement of motor and perceptual awareness when learning to reach with mirror reversed feedback

**DOI:** 10.64898/2026.07.30.741776

**Authors:** Sarvenaz Heirani Moghaddam, Gerome Aleandro Manson, Erin Krista Cressman

**Author notes:** <u>Corresponding Author</u>.

## Abstract

In mirror reversed (MR) learning, the magnitude and direction of the visuomotor distortion varies with target location. To date, implicit (i.e., unconscious) processes have not been implicated in learning to reach with an MR distortion, even when the distortion is small in magnitude. Across two experiments, we examined whether explicit processes (i.e., motor and perceptual awareness of reaching strategies) are engaged when learning to reach with a small (20°) MR distortion and whether this learning generalizes to novel targets. Learning to reach with an MR distortion was compared to learning to reach with a small visuomotor rotation (VR), in which cursor feedback was rotated 20° relative to hand motion at each target. Participants in the MR group engaged both motor and perceptual awareness and learning to reach with the MR distortion generalized to novel targets. Participants in the VR group also learned to reach with the VR distortion but they did not engage either motor or perceptual awareness and there was no evidence of generalization. Reaction times were longer for the MR group compared to the VR group, consistent with engagement of explicit processes. Together, these findings suggest that learning to reach with an MR distortion is supported by motor and perceptual awareness that generalize to novel targets.

## Introduction

Mirror reversed (MR) visuomotor distortions present cursor feedback reflected across the body midline (y-axis). As a result, movements to the right of the body midline result in cursor feedback displayed on the left side of the workspace, and vice versa (Telgen et al., 2014; Wang & Taylor, 2021; Wilterson & Taylor, 2021). Because the cursor distortion is dependent on how far the hand is from one’s body midline, the magnitude and direction of the distortion vary across the workspace, and hence are related to target location (Telgen et al., 2014; Wang & Taylor, 2021; Wilterson & Taylor, 2021). Previous research has indicated that participants do not display aftereffects (i.e., unconscious, implicit learning) after reaching with an MR distortion. That is, once the MR distortion is removed, participants immediately resume reaching directly to the target with minimal error (Telgen et al., 2014; Wang & Taylor, 2021; Wilterson & Taylor, 2021). The absence of aftereffects has been taken as evidence that implicit learning does not contribute to learning to reach with an MR distortion (Wang & Taylor, 2021; Wilterson & Taylor, 2021; Heirani Moghaddam, Cressman & Manson, submitted).

The absence of implicit learning when reaching with an MR distortion, regardless of distortion size (e.g., 20° to 90°; Wang & Taylor, 2021; Wilterson & Taylor, 2021; Heirani Moghaddam, Cressman & Manson, submitted), contrasts with learning to reach with a small visuomotor rotation (VR) distortion. When reaching with a VR distortion, visual feedback is rotated clockwise (CW) or counterclockwise (CCW) relative to one’s hand motion (Baraduc & Wolpert, 2002; Bastian, 2008; Ghahramani et al., 1996; Vetter et al., 1999). For example, a 20° CW VR distortion requires participants to reach 20° CCW of the target, regardless of the position of the target in the workspace (Cressman & Henriques, 2009; Krakauer, 2009; Krakauer et al., 2005). After learning to reach with a VR distortion, participants exhibit consistent aftereffects such that they continue to reach in the opposite direction of the rotation when the distortion is removed. (Krakauer et al., 1999; Shadmehr & Mussa-ivaldi, 1994). These aftereffects reflect implicit learning, which has been shown to be responsible for learning to reach with VR distortions of less than 30° (Modchalingam et al., 2019; Neville & Cressman, 2018; Werner et al., 2015).

While the engagement of implicit processes depend on the type of visuomotor distortion introduced (i.e., MR versus VR distortion), recent work indicates that explicit processes (i.e., conscious strategies) contribute to learning to reach with large MR distortions (Wang & Taylor, 2021; Wilterson & Taylor, 2021) and large VR distortions (Maresch et al., 2020; Neville & Cressman, 2018; Werner et al., 2015), where large distortions are typically defined as distortions greater than 30° (Heirani Moghaddam et al., 2021; Modchalingam et al., 2019). Efforts to characterize explicit learning when reaching with a VR distortion have employed motor and perceptual assessments of reaching strategies. Motor awareness reflects participants’ conscious monitoring and adjustment of their movements from trial to trial, thereby giving rise to awareness of the reaching strategies they employ (Johnson & Haggard, 2005; Werner et al., 2015). Perceptual awareness reflects participants’ conscious knowledge of the visuomotor distortion itself, independent of movement execution (Johnson & Haggard, 2005).

Within VR paradigms, motor awareness has been assessed using the process dissociation procedure (PDP; Heirani Moghaddam et al., 2021; Jacoby, 1991; Werner et al., 2015). The PDP requires participants to reach under two sets of instructions: (1) inclusion instructions, in which participants reach using any strategy they have learned, and (2) exclusion instructions, in which participants reach in the absence of any strategy they have learned. Explicit learning is quantified as the difference in reaching errors between inclusion and exclusion trials, while implicit learning is established based on errors in the exclusion trials. Perceptual awareness has been assessed by requiring participants to directly report their planned aiming direction by either (1) verbally indicating a number from an array of numbers surrounding the target that corresponds to their planned aiming direction (Heirani Moghaddam et al., 2021; Maresch et al., 2020; Taylor et al., 2014) or (2) orienting a line with respect to the target to indicate their planned aiming direction (line orienting task: LOT).

According to the action-perception framework, visual information for action versus perception is processed through distinct pathways: the dorsal stream supports visual processing for the control of action, whereas the ventral stream supports visual processing for perceptual judgments (Goodale & Milner, 2018, 1992; Milner, 2017; Milner & Goodale, 2008). Thus, given that the PDP and LOT assess motor versus perceptual awareness respectively, these assessments may engage partially distinct neural substrates. The PDP likely engages the dorsal stream, reflecting action-related awareness available during movement, whereas the LOT may more strongly engage the ventral stream, reflecting perception of a planned aiming direction. Consistent with this proposed dissociation, results from VR paradigms indicate that one’s motor awareness (PDP assessment) and perceptual awareness (LOT assessment) of reaching strategies reflect distinct yet complementary aspects of explicit learning (Heirani Moghaddam et al., 2021, 2022; Maresch et al., 2020, 2021). Specifically, these assessments yield differences with respect to the magnitude of explicit learning. Maresch et al. (2020) found that, regardless of assessment frequency, perceptual awareness was significantly greater than motor awareness. Further supporting this dissociation, motor awareness combined with implicit learning underestimates total learning, whereas perceptual awareness combined with implicit learning closely approximates total learning, particularly when visual feedback is available (Heirani Moghaddam et al., 2021; Maresch et al., 2020; Modchalingam et al., 2019; Taylor et al., 2014; Werner et al., 2015). Together, these findings suggest that motor and perceptual awareness capture distinct aspects of explicit learning.

Given the dissociation between motor and perceptual measures of explicit learning in VR paradigms, analogous differences in movement strategies may emerge when learning to reach with an MR distortion, depending on the assessment method. Studies have shown that participants engage perceptual awareness when learning to reach with a large MR distortion (Wang & Taylor, 2021; Wilterson & Taylor, 2021). However, it remains unclear whether perceptual awareness is also engaged when learning to reach with a small MR distortion (i.e., 20°) and whether motor awareness, as assessed via the PDP, is engaged. Thus, our understanding of the extent to which explicit processes contribute to learning to reach with an MR distortion and whether motor and perceptual awareness capture distinct components of these processes when an MR distortion is introduced, is currently limited.

In the present study we sought to characterize the explicit processes engaged when learning to reach with an MR distortion using both motor and perceptual assessments. Specifically, we assessed the extent to which motor awareness (PDP assessment) and perceptual awareness (LOT assessment) support learning to reach in an MR environment. We further compared explicit learning during reaching with an MR distortion to explicit learning during reaching with a VR distortion to determine if they engage similar explicit processes. Finally, we examined the generalization of explicit learning to novel targets. Generalization reflects the extent to which learning transfers to novel contexts, such as untrained target directions (Cressman & Henriques, 2015; Gastrock et al., 2024). In VR paradigms, generalization to novel targets is typically limited when implicit processes are engaged, but is greater when explicit processes are engaged (Cressman & Henriques, 2015; Heuer & Hegele, 2011). A recent study reported generalization of learning to reach with an MR distortion across effectors and across the mirroring axis when targets were presented on only one side of the workspace (Gastrock et al., 2024); however, how explicit learning supports this generalization remains unclear.

We hypothesized that learning to reach with an MR distortion would rely primarily on explicit processes, including both motor awareness (PDP assessment) and perceptual awareness (LOT assessment). We further expected motor awareness to be less accurate than perceptual awareness because motor awareness depends on additional stages of action selection and execution, whereas perceptual awareness reflects the intended movement plan prior to execution (Sober & Sabes, 2003, 2005). Finally, we expected motor and perceptual awareness following reaches with an MR distortion to demonstrate generalization to novel targets. In contrast, we did not expect motor and perceptual awareness to be engaged in learning to reach with a VR distortion. Understanding the distinct engagement of motor and perceptual awareness during learning to reach with small MR and VR distortions will provide insight into how these sensorimotor mappings are learned.

## Methods

### Participants

Eighty participants were recruited from Queen’s University across two Experiments. In Experiment 1, 40 participants aged 18–50 years (mean age: 22 ± 3.5 years), were randomly assigned to either the MR group (N = 20; F = 11) or the VR group (N = 20; F = 12). The VR group was further divided into two subgroups consisting of 10 participants each: VR clockwise (VR-CW) and VR counterclockwise (VR-CCW). In Experiment 2, an independent sample of 40 participants aged 19–34 years (mean age: 21 ± 3 years), was similarly assigned to the MR group (N = 20; F = 11) or the VR group (N = 20; F = 14), with VR-CW and VR-CCW subgroups consisting of 10 participants each.

The experimental protocol for both experiments was approved by the Queen’s University Health Sciences and Affiliated Teaching Hospitals Research Ethics Board (HSREB). Written informed consent was obtained from all participants, and participants were informed of their right to withdraw from the experiment at any time without consequence. Upon arrival at the laboratory, participants completed the Edinburgh Handedness Inventory (Experiment 1: mean score = 94.9, SD = 7.9; Experiment 2: mean score = 92.8, SD = 11.5), confirming that all participants were right-handed. Participants also completed a brief neurological questionnaire (adapted from Miles, 1930) to confirm the absence of neurological impairment.

### Experimental Apparatus and Procedures

The two Experiments followed a similar protocol to that used previously (see Heirani Moghaddam, Cressman & Manson, submitted). Participants performed slicing (shooting) movements with their right hand using the Kinarm Exoskeleton (Kinarm, Kingston, ON, Canada; Kasuga et al., 2022; Scott, 1999; Singh & Scott, 2003). The Kinarm was positioned adjacent to the experimenter’s computer workstation and consisted of a downward-facing computer monitor (120 Hz refresh rate) and a reflective surface located 20.5 cm beneath the monitor. The downward-facing monitor projected visual stimuli onto the reflective surface, such that cursor feedback appeared spatially aligned with the participant’s hand, which was 20.5 cm below the reflective surface. Participants were seated in a height-adjustable wheelchair, and their arms were placed in adjustable troughs on the exoskeleton, with the left arm remaining stationary throughout the experiment. The experimenter then adjusted the chair height and distance from the reflective surface to ensure comfortable viewing and reaching to the visual stimuli. Participants performed flexion, extension, abduction and adduction movements of the right arm in the horizontal plane. The right index finger’s position was displayed as a cursor (0.5 cm diameter white circle) and tracked at a sampling rate of 1000 Hz. To eliminate visual feedback of the right limb, a drape was secured around the participant’s neck with Velcro. The drape, along with the reflective surface, fully obscured the participant’s view of their right arm.

### Trials

Participants made reaching movements towards two targets positioned 10 cm from the home position (1 cm in diameter). The targets were blue circles, 0.75 cm in diameter, presented with equal probability. The y-axis (i.e., the mirroring axis) was aligned with a participant’s midline. The right target was positioned 10° to the right of the y-axis and the left target was positioned 10° to the left of the y-axis (Figure 1A). In Experiment 2, novel targets were also presented such that they were positioned 25° to the right and left of the y-axis (Figure 1B). The *target array* was defined as the circumference of an unseen circle centered on the home position, with a radius of 10 cm corresponding to the distance between the home position and target locations (Figure 1C and Figure 1D semicircular dotted line).

**Figure 1.**
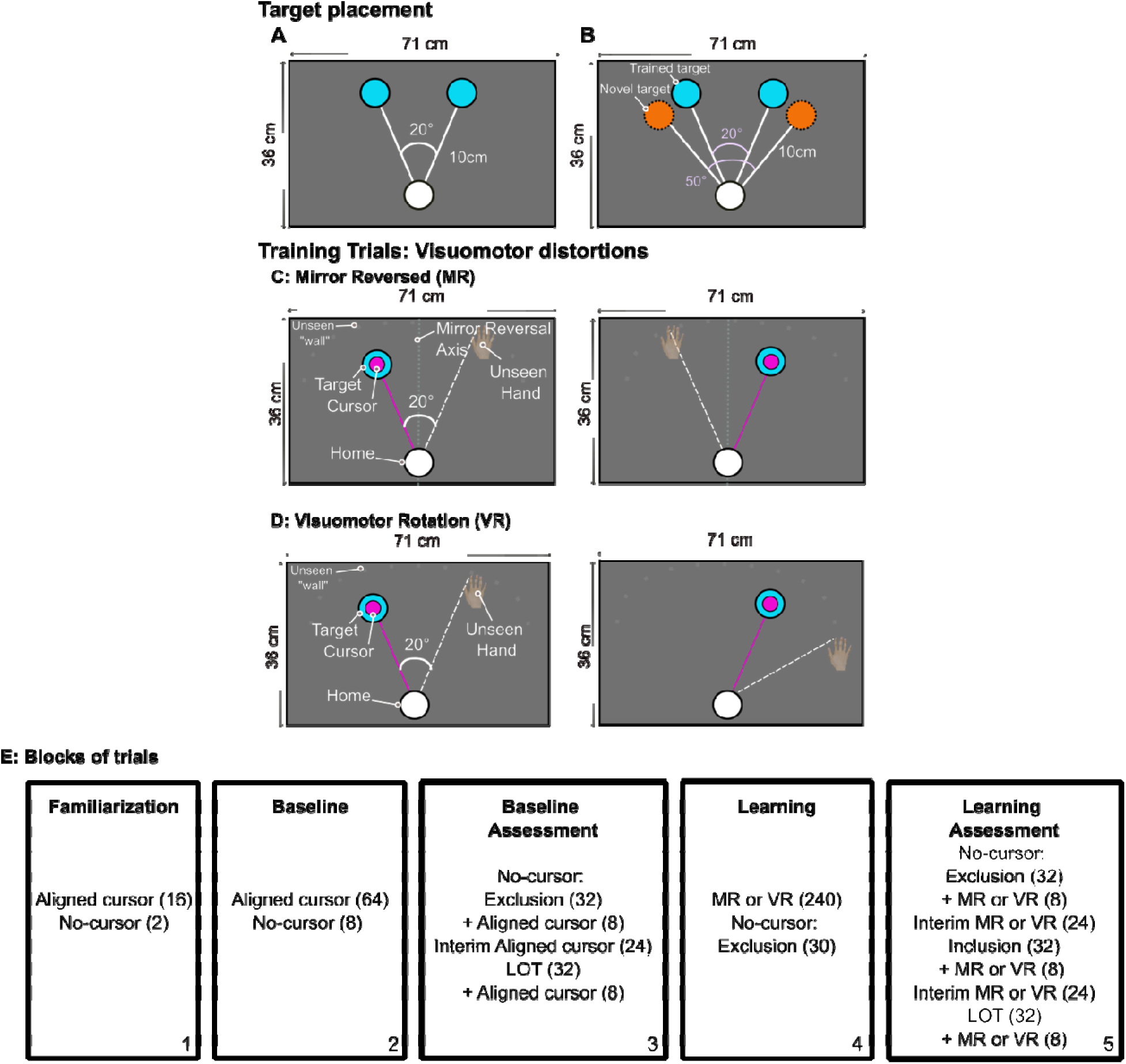
Target placement and visuomotor distortions. **A:** In both Experiments, the trained (familiar) targets were positioned 10° to the right and left of the body midline (y-axis). **B:** In Experiment 2, during the baseline and learning assessment blocks, novel (unfamiliar) targets appeared 25° to the right and left of the body midline, in addition to the trained targets. **C:** For the mirror reversed (MR) group, cursor motion was mirrored across the body midline (y-axis; dark grey dashed line in C) relative to the index finger motion. When the left (right) target was shown, the index finger had to move 20° to the right (left) of the target for the cursor to land on the target. **D:** For the visuomotor rotation (VR) group, cursor motion was rotated 20° clockwise (CW) or counterclockwise (CCW) relative to the index finger motion. For the CCW rotation shown in D, participants had to reach 20° to the right of the target for the cursor to land on both the right and left targets. **E:** Breakdown of blocks of trials and the number of trials (in parentheses) for each group in both Experiments. In Experiment 2, participants completed the same trial structure except that no-cursor trials were not performed in the baseline (block 2) and learning (block 4) blocks. In blocks 3 and 5, participants in Experiment 1 completed 16 trials to each target, whereas participants in Experiment 2 completed 8 trials to each of the four targets.

Each trial began with participants positioning their right index finger at the home position for 500 ms (see Figure 2). In Experiment 1, a yellow target was presented at the same time as the home position. Following a variable delay of 300-700 ms, the target turned blue (go signal). In Experiment 2, following the presentation of the home position and a variable delay of 300-700 ms, a blue target appeared. Participants performed rapid “slicing” movements through the target, consistent with ballistic shooting movements as described by Maksimovic et al. (2020). Movements were terminated by a soft mechanical wall (spring constant: 150 N/m; not seen by the participant), that was placed 2 cm beyond the *target array* to encourage natural movement dynamics and promote rapid execution.

**Figure 2.**
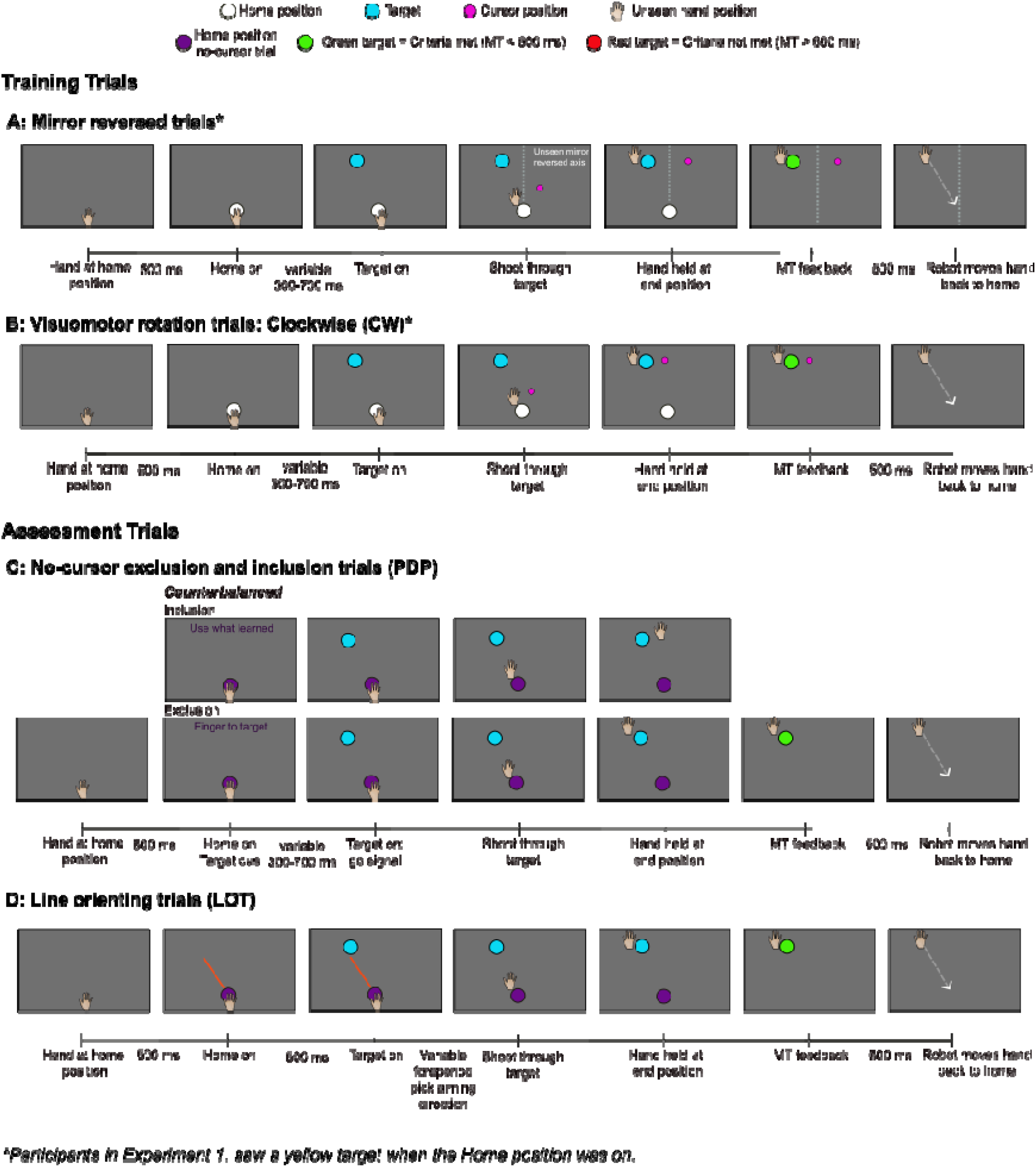
Timeline for trials completed. The hand and index finger were hidden from participants’ view. **Training Trials. A:** Mirror reversed (MR) trials: The cursor’s trajectory was mirrored across the y-axis relative to the trajectory of the index finger and was visible throughout the reach until the finger crossed the target array. **B:** Visuomotor rotation (VR) trials: The cursor’s trajectory was rotated 20° CW or 20° CCW relative to the trajectory of the index finger and was visible throughout the reach until the finger crossed the target array. **Assessment Trials. C:** No-cursor PDP assessment trials: Inclusion (top row) and Exclusion (bottom row) trials. **D:** Line orienting trials (LOT): Participants first aligned the red line to indicate where they thought their finger must aim to get the cursor to the target and then reached to the target with no cursor feedback.

Cursor feedback of the index finger’s position was available until the finger crossed the target array, after which the cursor disappeared. Movement onset was defined online as the time when the center of the cursor moved 0.5 cm outwards from the center of the home position and the cursor’s velocity rose above 0.03 m/s and remained above that threshold for at least 9 ms. Movement end was recorded when the participant’s finger crossed the target array, and movement time (MT) was calculated as the time interval between movement onset and movement end.

Participants received MT feedback at the end of each trial via a target color change: a green target indicated the goal MT of less than 600 ms had been achieved, while a red target signaled the actual MT was longer than the goal MT (i.e., MT > 600 ms). This feedback was provided as the hand remained stationary at the movement endpoint for 500 ms. Following this delay, the Kinarm moved the finger back to the home position along a straight trajectory over a duration of 1000 ms. This return movement occurred in the absence of visual feedback.

### Types of Trials Training trials

Participants performed shooting movements under one of three feedback conditions: aligned cursor trials, mirror reversed (MR) trials, and visuomotor rotation (VR) trials. In aligned training trials, a white cursor (0.5 cm in diameter) accurately represented the index finger’s position, providing veridical visual feedback. Aligned training trials were completed in the familiarization, baseline and baseline assessment blocks (Figure 1E, blocks 1-3). In MR trials, the cursor trajectory was mirrored across the y-axis relative to actual hand motion (Figure 1C & Figure 2A; MR group). In VR trials, the cursor trajectory was rotated either 20° CW or CCW relative to actual hand motion (Figure 1D & Figure 2B; VR-CW and VR-CCW groups). Participants were not informed about the nature of the visuomotor distortions. MR and VR training trials were completed during the learning and learning assessment blocks (Figure 1E, blocks 4 and 5).

### Assessment trials

The timing of events in assessment trials was similar to that of the training trials (see Figure 2). All assessment trials were completed in the absence of cursor feedback, designed to isolate implicit and explicit learning. The absence of cursor feedback in the assessment trials was cued with a change in colour of the home position, such that on these trials the colour of the home position was purple (see Figure 2C and D).

### PDP: Exclusion and Inclusion Trials – Motor Awareness

Participants completed these no-cursor trials under exclusion and inclusion instructions. Figure 1E (blocks 3 and 5) depicts the number of PDP exclusion and inclusion trials completed and Figure 2C depicts the visual cues presented during these trials. The exclusion trials were used to establish implicit learning in accordance with the Process Dissociation Procedure (PDP; adapted from Werner et al., 2015). For these trials, the words “Finger to Target” appeared above the target and participants were instructed:

> *“You are now going to reach when you cannot see your* index finger*, as there will be no-cursor on the screen. Do not use anything you may have learned for these trials to get the cursor to the target. Instead, aim so that your* index finger *goes straight through the target as you did during baseline reaches.”*

For the no-cursor inclusion trials, participants were given these instructions:

> *“You are now going to reach when you cannot see your index finger, as there will be no-cursor on the screen. For these trials, use anything you have learned during training to get the cursor to the target. In other words, aim so that the cursor would have gone straight to the target, as in the training trials you just completed.”*

When participants were completing the inclusion trials, the words “Use what learned” appeared above the target. These trials were completed in block 5 of both Experiments.

### LOT: Line Orienting Trials – Perceptual Awareness

Figure 1E (blocks 3 and 5) depicts the number of LOT trials completed and Figure 2D depicts the visual cues presented during these trials. A trial began with a red line (length = 10 cm, diameter = 0.25 cm) presented at the home position, randomly positioned in the first or second quadrant of the Cartesian plane. The red line then rotated from the right side to the left or the left side to the right (i.e., 180°) over 5000 ms. Participants said ‘stop’ to indicate when the experimenter should press a button to stop the movement of the line. The position of the line indicated the direction the participant planned to reach to get the cursor to the target. Once the line had stopped, participants instructed the experimenter to move the line left or right by 1° increments to adjust its position until they were confident in the orientation chosen. Once participants chose their aiming direction, the line disappeared, and participants reached to the target with no-cursor feedback. In Experiment 1, the target was yellow while participants positioned the red line and then the target turned blue to indicate they could begin their reach. In Experiment 2, the target was blue from the start of a trial. Participants were to initiate their movements when the orienting line disappeared.

The order in which the assessment trials were completed in the baseline and learning assessment blocks was counterbalanced across participants. For example, if one participant completed the learning assessment block in the following order: LOT trials followed by PDP exclusion and inclusion trials, then the next participant completed the block in a different order: PDP exclusion and inclusion trials, followed by the LOT trials. In Experiment 2, exclusion and inclusion trials, and the LOT trials were completed to 4 targets: the two trained targets and the two novel targets (left and right novel targets; Figure 1B). In the baseline assessment block and the learning assessment block, participants completed 24 interim trials with cursor feedback between exclusion, inclusion and LOT trials to maintain any motor learning.

### Data Analyses

All reaching trials were analyzed using custom-written MATLAB scripts (R2024a; The MathWorks, Inc.). The primary dependent variable was angular error at movement endpoint (AE), defined as the angular difference between a reference vector (home position to target) and the reach vector (home position to finger position at the target array). Other variables of interest included reaction time (RT) and movement time (MT). RT was calculated as the time from target onset to movement onset (defined as when the cursor moved 0.5 cm away from the home position and velocity exceeded 0.03 m/s for at least 9 ms). MT was computed as the time from movement onset to movement end (when the index finger crossed the target array).

### Outlier Detection

Five variables were examined to establish outlier trials: Start coordinates (Start X and Start Y), AE, RT and MT. A trial was discarded if Start X, Start Y, or AE exceeded 3 standard deviations (SD) from a participant’s mean value for the same trial type within a given block of trials (i.e., baseline, baseline assessment, learning, or learning assessment). For example, each aligned training trial was compared to the mean of aligned training trials within the same block, while no-cursor trials were assessed relative to their block-specific mean. With respect to RT and MT, any trial with RT or MT shorter than 100 ms was excluded, as well as trials in which MT was longer than 600 ms. In Experiment 1, one MR participant was excluded due to too many missing trials. In Experiment 2, one MR participant was excluded as they did not complete the full set of trials. Across both Experiments, 38,067 trials were collected, of which 2,555 trials (6.7%) were excluded from analyses. Exclusions per participant ranged from 0–18%. Importantly, including these trials did not alter statistical outcomes or the main effects reported below.

### Learning

To evaluate learning, AE was compared between baseline and learning blocks for each experiment. Each block was divided into early and late trials (first 24 trials versus last 24 trials across the left and right targets). Learning was established by comparing mean AE of the late learning trials to the mean AE of the late baseline trials. A participant was classified as having learned to reach with the distortion if late learning AE exceeded late baseline AE by more than 3 SD for each target (Heirani Moghaddam et al., 2025). This change in AE had to be in the direction that compensated for the specific distortion at both targets. In both Experiments, all VR participants met this criterion. However, 6 MR participants in Experiment 1 and 7 participants in Experiment 2 did not meet this threshold and were excluded from further analyses. This resulted in the inclusion of data from 13 MR participants in Experiment 1 and 12 MR participants in Experiment 2 in the main analyses. Data related to the excluded MR participants (MR non-learners, MR-NL) are presented in the Supplementary File. AEs in the learning block were then normalized by subtracting baseline AEs for the same target and time point (early or late trials), yielding normalized learning values for group comparisons.

The absolute values of normalized AEs were compared across groups in a 3 group (VR- CW, VR-CCW, MR) x 2 target position (right, left) x 2 time (early, late) mixed analysis of variance (ANOVA) with repeated measures (RM) on the last two factors. AE variability, RT and MT at the same time points were also analyzed in a 3 group x 2 target position x 2 time mixed ANOVA with RM on the last two factors. To note, VR participants were kept within their distinct subgroups for analyses (VR-CW or VR-CCW) to maintain similar sample sizes across the three groups.

### Implicit and Explicit Learning

Implicit and explicit learning were established via the PDP and LOT trials. All values were normalized such that positive values indicated a deviation in the expected direction of learning (i.e., + 20°).

### Implicit Learning: PDP

The implicit index was calculated for each target within the baseline and learning assessment blocks. The implicit index (I_Ind_) was defined as the mean AE of the no-cursor exclusion trials. For example:

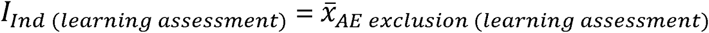

This index was then used to calculate implicit learning according to the following formula:

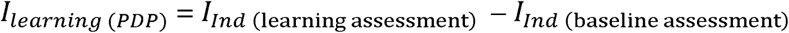

### Explicit Learning: Motor Awareness (PDP)

The explicit index (E_Ind_) in the learning assessment block and motor awareness (E_Motor Awareness (PDP)_) was established according to the following formulas:

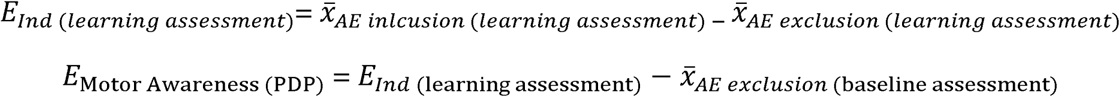

### Implicit Learning: LOT

Following each line orientation trial, participants reached to the target. The difference between this reach vector (between the home position and movement end) and the oriented line vector provided an index of implicit learning according to the following formulas:

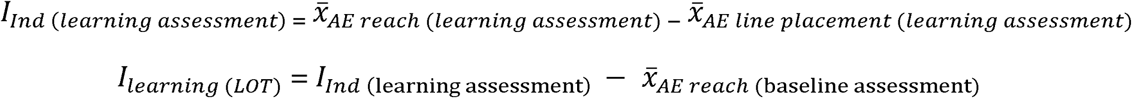

### Explicit Learning: Perceptual Awareness (LOT)

Participants’ perceptual awareness was defined as the angular difference between the oriented line vector and the reference vector from the home position to the target (see Figure 2D). This value was normalized to baseline for each target:

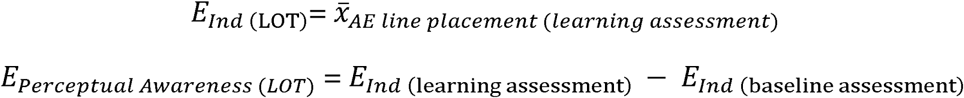

In Experiment 1, all implicit and explicit measures were compared across groups within a 3 group (VR-CW, VR-CCW, MR) × 2 target position (right, left) mixed ANOVA with RM on the last factor. In Experiment 2, all implicit and explicit measures were analyzed in a 3 group (VR- CW, VR-CCW, MR) x 2 target position (right, left) x 2 target type (trained, novel) mixed ANOVA with RM on the last two factors. We compared the magnitude of implicit and explicit learning established via the PDP and LOT trials in each group to zero using one-sample *t*-tests to determine whether these processes were present following learning to reach with an MR or VR distortion. To examine the relationship across assessments, Pearson correlations were conducted between motor awareness (PDP) and perceptual awareness (LOT) for both experiments.

### Explicit Learning in MR

In addition, we quantified absolute errors in explicit learning for both motor and perceptual awareness as assessed via the PDP and LOT, respectively. This error was defined as the absolute angular deviation between a participant’s awareness estimate and the reference angle. For targets in Experiments 1 and trained targets in Experiment 2, the reference angle corresponded to ±20° (−20° for right targets; +20° for left targets). For novel targets in Experiment 2, the reference angle corresponded to ±50° (−50° for right targets; +50° for left targets). Errors were computed as the absolute difference between the participant’s awareness estimate (PDP or LOT) and the corresponding reference angle:

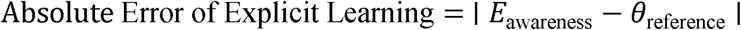

where *E*_awareness_ denotes the participant’s awareness (motor or perceptual) and *θ*_reference_ denotes the specific reference angle. Because the same trained targets were used in Experiment 1 and Experiment 2, absolute error data from both Experiments were included in paired-samples *t*-tests to examine differences between assessment methods (PDP vs LOT). The same analysis was conducted separately for the novel targets in Experiment 2.

All statistical analyses were conducted using JASP, RStudio, and Microsoft Excel. For RM ANOVA, sphericity was assessed using Mauchly’s test and no violations were observed. The alpha level for significance was set at *p* < 0.05 and post hoc comparisons were performed with Bonferroni correction where appropriate. Effect sizes are reported as partial eta squared (η*_p_²*) or Cohen’s d (*d*). In addition, coefficients of determination (*R^2^*), and slopes for linear regressions are reported to describe the relationship between the variables of interest where applicable. For clarity, results focus on the highest-order significant interactions and lower-order effects are reported only when relevant.

## Results

### Learning Experiment 1

For learning, ANOVA revealed significant main effects of group (*F*(2,30) = 12.502, *p* < 0.001, η*_p_²* = 0.455), and time (*F*(1,30) = 96.512, *p* < 0.001, η*_p_²* = 0.763), and a significant interaction between group x time (*F*(2,30) = 22.563, *p* < 0.001, η*_p_²* = 0.601). Post hoc analyses revealed that early AEs were significantly smaller than late AEs for the MR group and the VR- CCW group, indicating greater learning at late relative to early trials (all *p* < 0.05). In addition, early AEs for the MR group were significantly smaller than early AEs in the VR-CW and VR- CCW groups (all *p* < 0.001). No significant difference was observed for the VR-CW group between early and late trials (*p* = 0.204). Importantly, late AEs were not significantly different between groups (all *p* = 1.000; Figure 3A and Figure 3B).

**Figure 3.**
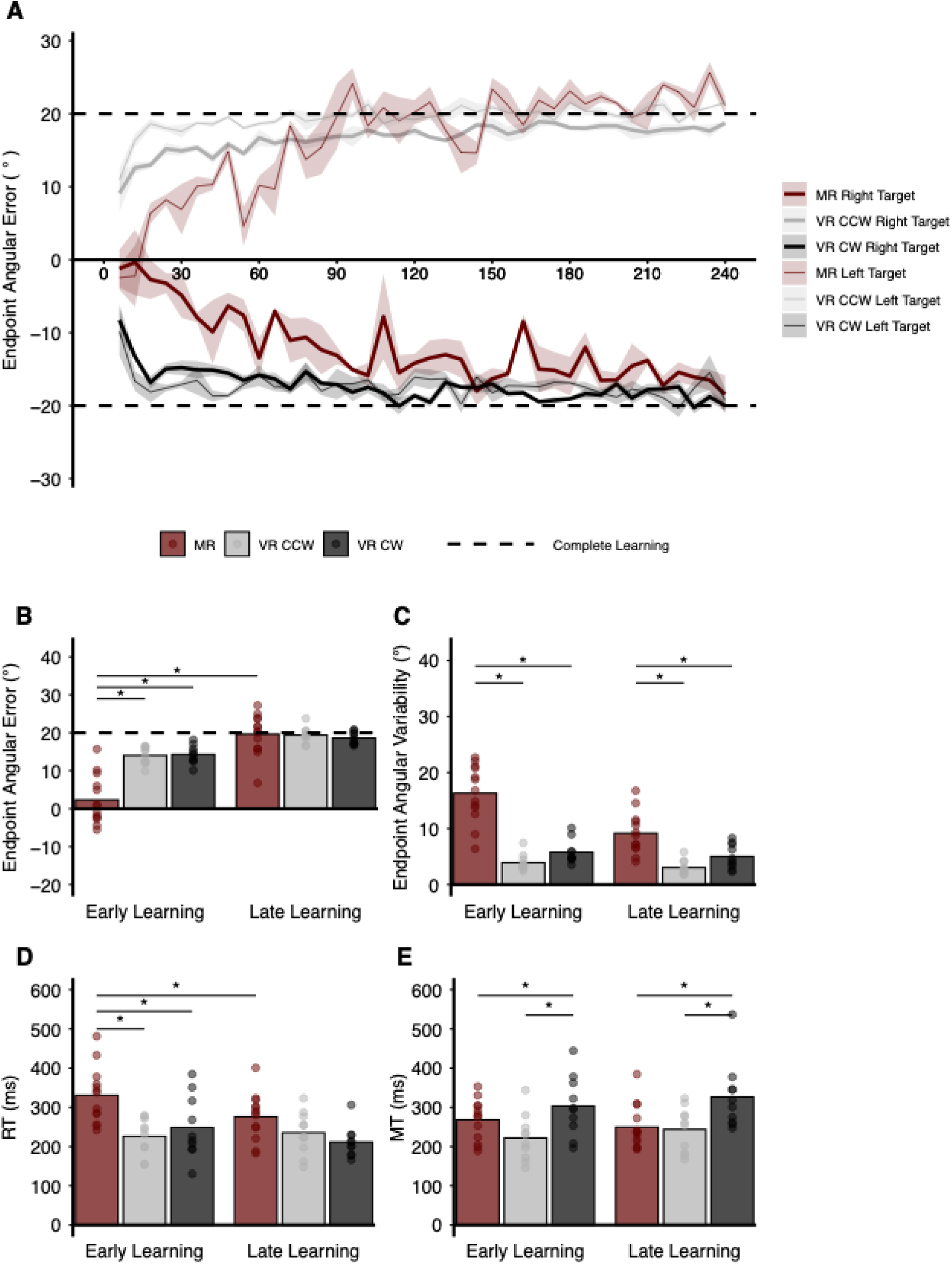
Learning performance across groups in Experiment 1. **A:** Learning curves for each group across trials. Each point represents the mean endpoint angular error (AE, in degrees) of three consecutive trials in the learning block. Bold solid lines reflect reaches toward the right target, while thin solid lines indicate reaches toward the left target. Shaded regions represent the standard error of the mean (SEM). Dashed lines indicate the expected reach adjustments for complete learning: y = −20° for the right target, and y = 20° for the left target. Group colors: MR (maroon), VR-CCW (gray), and VR-CW (black). **B:** Absolute AE, **C:** Variability in AE (standard deviation, SD), **D:** Reaction time (RT), and **E:** Movement time (MT) data for each group during early and late learning. Individual participant data are represented as circles. Asterisks denote significant differences between groups at the same time, or within a group across times (*p* ≤ 0.05).

Although mean late AEs were similar across groups, AE variability differed significantly across groups (*F*(2,30) = 45.809, *p* < 0.001, η*_p_²* = 0.753), and across time (*F*(1,30) = 18.625, *p* < 0.001, η*_p_²* = 0.383). ANOVA further revealed a significant interaction between group x time (*F*(2,30) = 10.495, *p* < 0.001, η*_p_²* = 0.412). Post hoc analyses indicated that within the MR group, early AEs were significantly more variable compared to late AEs (*p* < 0.001). AE variability was also significantly larger for the MR group compared to the VR-CW group and the VR-CCW group across early and late learning trials (all *p* < 0.05; Figure 3C).

For RT, ANOVA revealed significant main effects of group (*F*(2,30) = 7.422, *p* = 0.002, η*_p_²* = 0.331), and time (*F*(1,30) = 7.084, *p* = 0.012, η*_p_²* = 0.191), and a significant interaction between group x time (*F*(2,30) = 3.378, *p* = 0.048, η*_p_²* = 0.184). Post hoc analyses indicated that RT on early learning trials for the MR group was significantly longer than RT on late learning trials (*p* = 0.035). Early RT for the MR group was also significantly longer than early RT for the VR-CCW group (*p* = 0.008). By late learning, RTs did not significantly differ across groups (all *p* = 1.000; Figure 3D).

Mean MTs remained within the prescribed MT criteria for all groups, ranging from 218 ms to 332 ms throughout the learning block. Analysis of MT revealed significant main effects of group (*F*(2,30) = 4.879, *p* = 0.015, η*_p_²* = 0.245), and target position (*F*(1,30) = 4.419, *p* = 0.044, η*_p_²* = 0.128), and significant interactions between target position × time (*F*(1,30) = 19.764, *p* < 0.001, η*_p_²* = 0.397) and group x target position x time (*F*(2,30) = 6.364, *p* < 0.001, η*_p_²* = 0.298). Post hoc analyses revealed that for the MR group, early MTs toward the right target were significantly shorter than early MTs towards the left target (*p* = 0.001). As well for the MR group, early MTs towards the left target were significantly longer than late MTs towards the same target (*p* = 0.017). Finally, the VR-CCW group had significantly shorter MTs compared to the VR-CW group (*p* = 0.014; Figure 3E).

### Experiment 2

The results of Experiment 2 are similar to Experiment 1 and are reported in detail in the Supplementary File. In brief, ANOVA revealed a significant main effect of time (*F*(1,29) = 25.819, *p* < 0.001, η*_p_²* = 0.471), and a significant interaction between group x time x target position (*F*(2,29) = 5.324, *p* = 0.011, η*_p_²* = 0.269) with respect to mean AE. Post hoc analyses revealed that early AEs were similar across groups (*p* > 0.05), however, for the MR group, early AEs were significantly different from their late AEs and from late AEs in the VR-CW group and the VR-CCW group (all *p* < 0.05). Importantly, all participants learned their respective distortions to a similar extent by late learning such that late AEs toward the right and left targets did not significantly differ between groups (all *p* > 0.05; Figure S1A and S1B). AE variability was significantly larger for the MR group compared to the VR-CW group and the VR-CCW group across early and late trials (all *p* < 0.05; Figure S1C), and RTs were significantly longer for the MR participants in early and late learning trials compared to the VR-CW and VR-CCW groups (all *p* < 0.001; Figure S1D). In terms of MT, all groups completed reaches well within the 600 ms MT criterion across the learning block. The VR-CW group did reach with significantly longer MTs compared to the MR and VR-CCW groups in early and late learning (all *p* < 0.05; Figure S1E).

### Implicit Learning: PDP

In Experiment 1, implicit learning (PDP) differed significantly between groups (*F*(2,30) = 42.725, *p* < 0.001, η*_p_²* = 0.740), with the MR group showing significantly less implicit learning (M = –2.8° ± 5.8°) than both the VR-CW group (M = 12.5° ± 4.6°) and the VR-CCW group (M = 12.3° ± 6.7°; Figure 4A). Comparison of the magnitude of implicit learning for the MR group to zero revealed a significant difference in the opposite direction of expected learning (*t*(12) = - 2.99, *p* = 0.011, *d* = 0.321).

**Figure 4.**
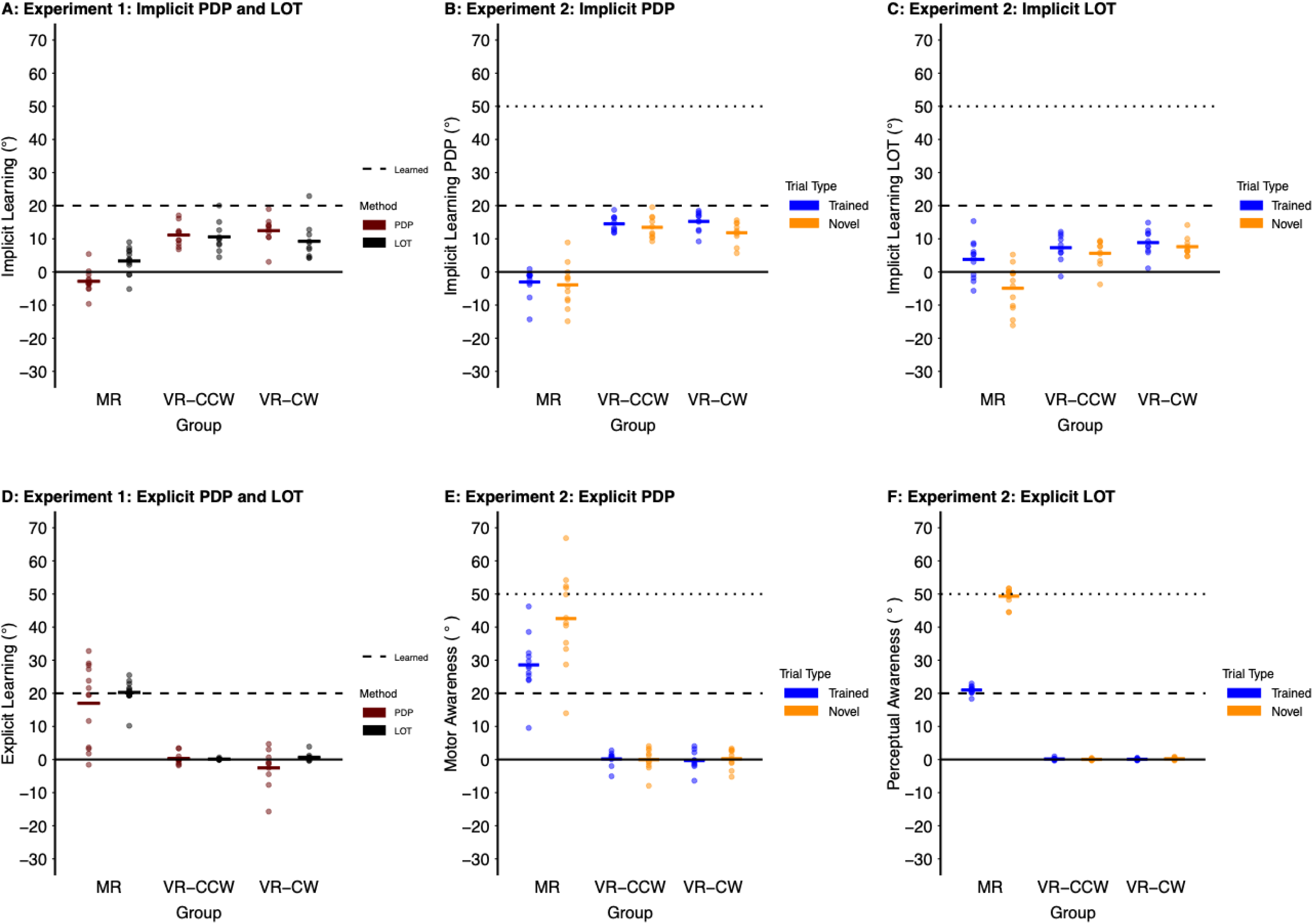
Implicit learning (A, B and C) and explicit learning (D, E and F) across Experiments, groups and assessment methods. All figures show group-level (dark, bold line) and individual implicit and explicit learning (circles) established via the Process Dissociation Procedure (PDP) and the Line Orienting task (LOT). **A:** Experiment 1 implicit learning as established via the PDP (maroon) and LOT (black) trials. **B:** Implicit learning established via the PDP in Experiment 2 to trained (blue lines and dots) and novel (yellow lines and dots) targets. **C:** Implicit learning established via the LOT in Experiment 2 to trained (blue lines and dots) and novel (yellow lines and dots) targets. **D:** Experiment 1 explicit learning established via the PDP (maroon; Motor Awareness) and LOT (black; Perceptual Awareness). **E:** Explicit learning established via the PDP (Motor Awareness) in Experiment 2 to trained (blue lines and dots) and novel (orange lines and dots) targets. **F:** Explicit learning established via the LOT (Perceptual Awareness) in Experiment 2 to trained (blue lines and dots) and novel (orange lines and dots) targets. The thick dashed line indicates complete learning for the trained target and the thin dashed line indicates complete learning for the novel target.

Experiment 2 revealed similar findings with respect to implicit learning. We found significant main effects of group (*F*(2,29) = 80.687, *p* < 0.001, η*_p_²* = 0.848), and target type (*F*(1,29) = 8.384, *p* = 0.007, η*_p_²* = 0.224), and a significant interaction between group x target type (*F*(2,29) = 3.405, *p* = 0.047, η*_p_²* = 0.190). Post hoc analyses revealed that, like Experiment 1, implicit learning for the MR group established at the trained and novel targets was significantly smaller compared to both the VR-CW group and the VR-CCW group (both *p* < 0.001), and in the opposite direction to expected learning (Figure 4B). No significant differences were found between the trained and novel targets in the VR-CW group and the VR-CCW group, with the exception that for the VR-CW group implicit learning was significantly reduced at the right novel target compared to the right trained target (right; *p* = 0.006).

### Implicit Learning: LOT

In terms of implicit learning (LOT) in Experiment 1, ANOVA revealed a significant main effect of group (*F*(2,30) = 8.211, *p* = 0.001, η*_p_²* = 0.354), with the MR group (*M* = 3.3° ± 5.6°) demonstrating significantly less implicit learning than the VR-CW group (*M* = 8.3° ± 6.9°; *p* = 0.014) and the VR-CCW group (*M* = 10.6° ± 4.5°; *p* = 0.002; Figure 4A). For the MR group, implicit learning established via the LOT differed significantly from zero (*M* = 3.3° ± 3.9°; *t*(12) = 3.103, *p* = 0.009, *d* = 0.325).

In Experiment 2, ANOVA revealed a significant main effect of group (*F*(2,29) = 10.940, *p* < 0.001, η*_p_²* = 0.430), and target type (*F*(1,29) = 58.817, *p* < 0.001, η*_p_²* = 0.670), and a significant interaction between group x target type (*F*(2,29) = 24.343, *p* < 0.001, η*_p_²* = 0.627). Post hoc analyses revealed that implicit learning established at the trained targets was similar across all groups (MR group: *M* = 3.8° ± 7.2°; VR-CW group: *M* = 8.8° ± 4.6°; VR-CCW group: *M* = 7.3° ± 4.5°; all *p* > 0.05). However, implicit learning established at the novel targets was significantly smaller for the MR group than both the VR-CW group and the VR-CCW group (MR group: *M* = -4.9° ± 8.8°; VR-CW group: *M* = 7.6° ± 3.6°; VR-CCW group: *M* = 5.6° ± 4.6°; all *p* < 0.05; Figure 4C). For the VR-CW group and the VR-CCW group, implicit learning did not differ between trained and novel targets (post hoc comparisons, all *p* > 0.05). For the MR group, one-sample *t*-tests revealed that implicit learning towards trained targets (*t*(11) = 2.235, *p* = 0.047, *d* = 0.645) and novel targets (*t*(11) = -2.478, *p* = 0.031, *d* = -0.715) significantly differed from zero. Specifically, for the MR group, implicit learning was in the direction of learning for trained targets, but in the opposite direction of expected learning for novel targets.

### Explicit Learning: Motor Awareness (PDP)

In Experiment 1, ANOVA revealed a significant main effect of group (*F*(2,30) = 15.980, *p* < 0.001, η*_p_²* = 0.504), with the MR group demonstrating significantly greater motor awareness (PDP) than both the VR-CW group and VR-CCW group (all *p* < 0.001; see Figure 4D & **Error! Reference source not found.**). As well, motor awareness in the VR-CW group (*t*(9) = -0.292, *p* = 0.777, *d* = -0.092) and the VR-CCW group did not significantly differ from zero (*t*(9) = 0.558, *p* = 0.591, *d* = 0.176).

In Experiment 2, ANOVA revealed significant main effects of group (*F*(2,29) = 91.556, *p* < 0.001, η*_p_²* = 0.863), and target type (*F*(1,29) = 38.555, *p* < 0.001, η*_p_²* = 0.571), and a significant interaction between group x target type (*F*(2,29) = 38.280, *p* < 0.001, η*_p_²* = 0.725). Post hoc comparisons indicated that motor awareness was significantly greater for the MR group compared to both the VR-CW group and VR-CCW group (both *p* < 0.001). As well, for the MR group, motor awareness was significantly larger at the novel targets compared to the trained targets (all *p* < 0.001; see Figure 4E & **Error! Reference source not found.**). Motor awareness in the VR-CW group (trained target: *t*(9) = -0.320, *p* = 0.756, *d* = -0.101; novel target: *t*(9) = 0.263, *p* = 0.799, *d* = 0.083) and the VR-CCW group (trained target: *t*(9) = 0.245, *p* = 0.812, *d* = 0.078; novel target: *t*(9) = -0.038, *p* = 0.971, *d* = -0.012) did not significantly differ from zero.

### Explicit Learning: Perceptual Awareness (LOT)

For Experiment 1, ANOVA revealed significant main effects of group *(F*(2,30) = 277.410*, p* < 0.001, η*_p_²* = 0.949), and target position (*F*(1,30) = 8.007*, p* = 0.008, η*_p_²* = 0.211), and a significant interaction between group x target position (*F*(2,30) = 3.673*, p* = 0.037, η*_p_²* = 0.197). Post hoc analyses indicated that the MR group exhibited significantly greater perceptual awareness at the right and left targets compared to the VR-CW group and the VR-CCW group (both *p* < 0.001), while the VR-CW group and the VR-CCW group did not significantly differ from each other (*p* = 1.000). For the MR group, perceptual awareness at the right target (*M* = 19.3° ± 3.5°) was significantly lower than the left target (*M* = 21.3° ± 4.1°; *p* = 0.004). Perceptual awareness in the VR-CW group (*t*(9) = 0.544, *p* = 0.600, *d* = 0.341) and the VR-CCW group did not significantly differ from zero (*t*(9) = –1.017, *p* = 0.336, *d* = 0.340; see Figure 4D & **Error! Reference source not found.**).

In Experiment 2, ANOVA revealed significant main effects of group (*F*(2,29) = 6334.932, *p* < 0.001, η*_p_²* = 0.998), and target type (*F*(1,29) = 957.798, *p* < 0.001, η*_p_²* = 0.971), and a significant interaction between group x target type (*F*(2,29) = 1010.690, *p* < 0.001, η*_p_²* = 0.986). Post hoc comparisons indicated that for the MR group, perceptual awareness established at the novel targets was significantly larger compared to the trained targets (*p* < 0.001; see Figure 4F & **Error! Reference source not found.**). As well, perceptual awareness in the VR-CW group (trained target: *t*(9) = 1.352, *p* = 0.209, *d* = 0.427; novel target: *t*(9) = 0.569, *p* = 0.583, *d* = 0.180) and the VR-CCW group (trained target: *t*(9) = 0.809, *p* = 0.439, *d* = 0.256; novel target: *t*(9) = 2.052, *p* = 0.070, *d* = 0.649) did not significantly differ from zero.

**Table 1.**
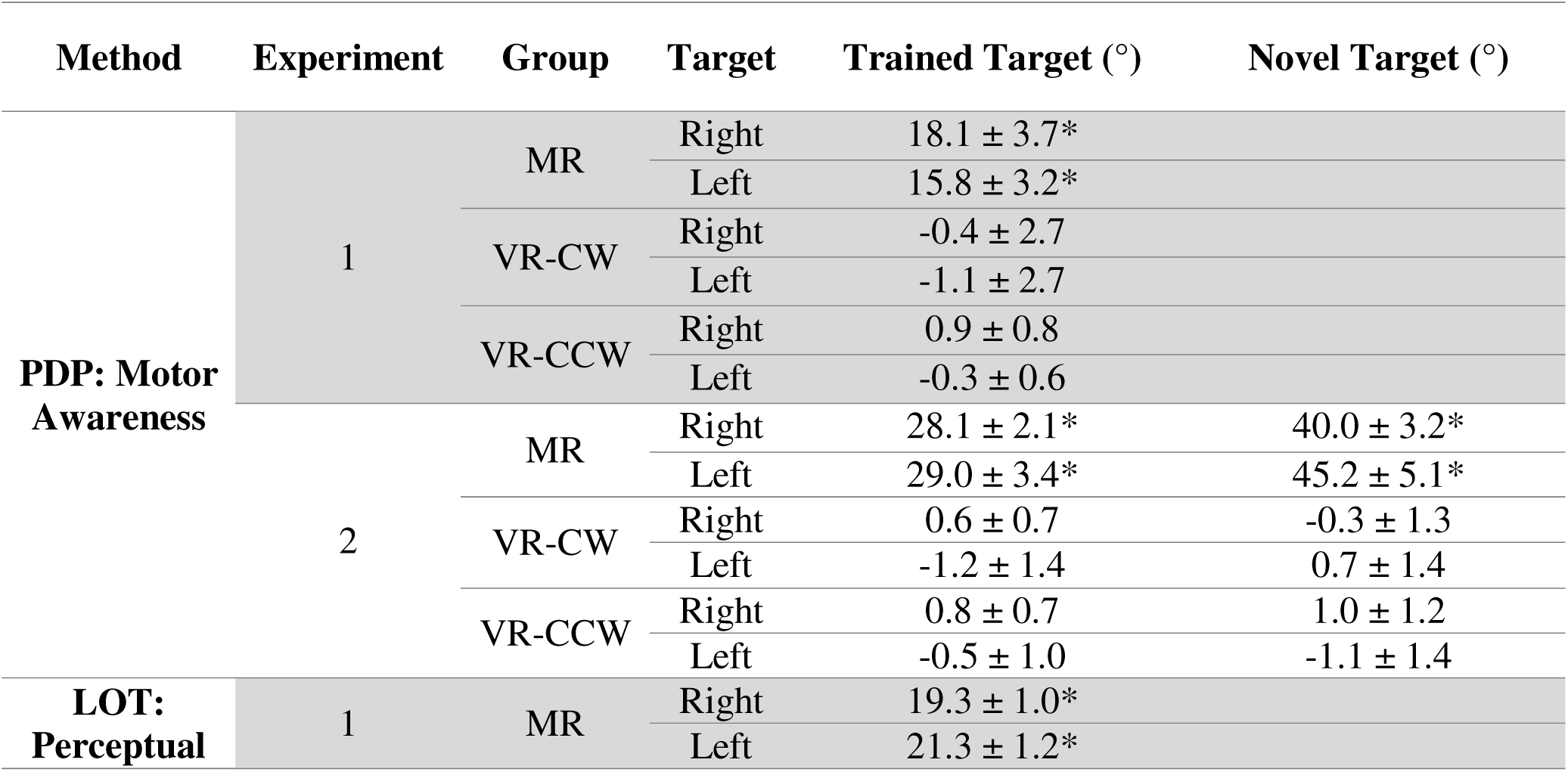
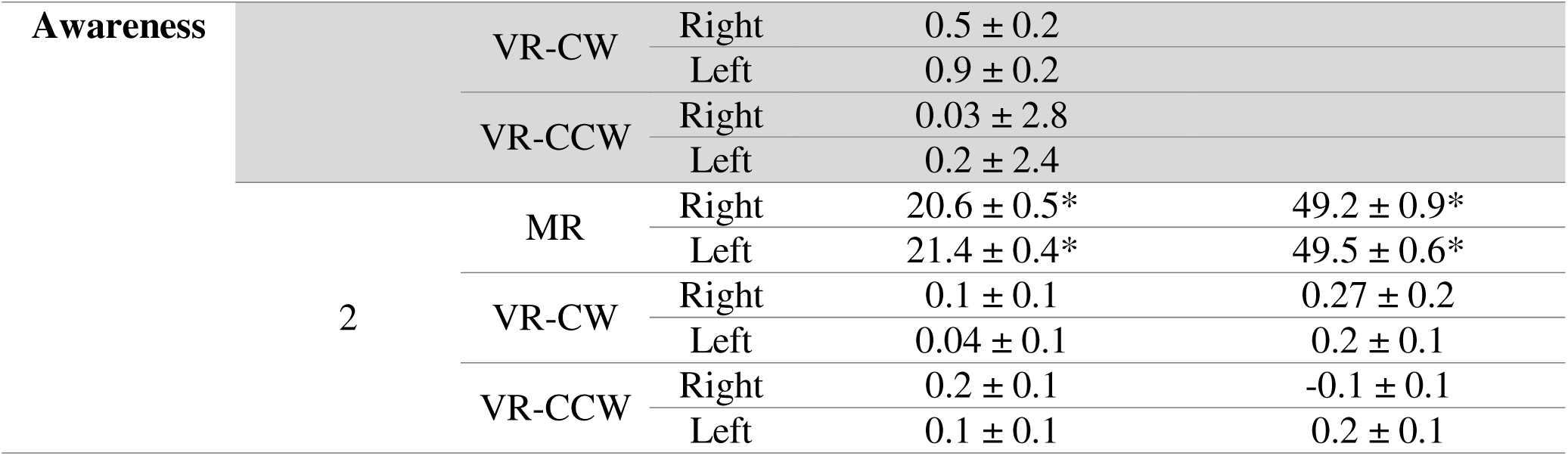
Post hoc comparisons of explicit learning (mean ± SD) as assessed via the Process Dissociation Procedure (PDP; motor awareness) and Line Orienting Task (LOT; perceptual awareness) at the trained (Experiments 1 and 2) and novel targets (Experiment 2) compared to zero. *Asterisks indicate a significant difference from zero.

### Explicit Learning in MR

Figure 5A, Figure 5B and Figure 5C illustrate absolute errors in explicit learning (motor and perceptual awareness) for the MR group at both trained and novel targets. Participants demonstrated significantly greater absolute errors on the LOT trials compared to the PDP trials, regardless of target type: trained targets: *t*(24) = 5.988, *p* < 0.001, *d* = 1.198 (Experiments 1 and 2); novel targets: *t*(11) = 4.562, *p* < 0.001, *d* = 1.317 (Experiment 2). Furthermore, in Experiment 2, absolute errors in explicit learning did not significantly differ between the trained and novel targets for the PDP (*t*(24) = -0.957, *p* = 0.359, *d* = -0.276) or the LOT (*t*(11) = 0.843, *p* = 0.417, *d* = 0.243), suggesting that participants were able to maintain similar absolute errors when novel targets were displayed.

**Figure 5.**
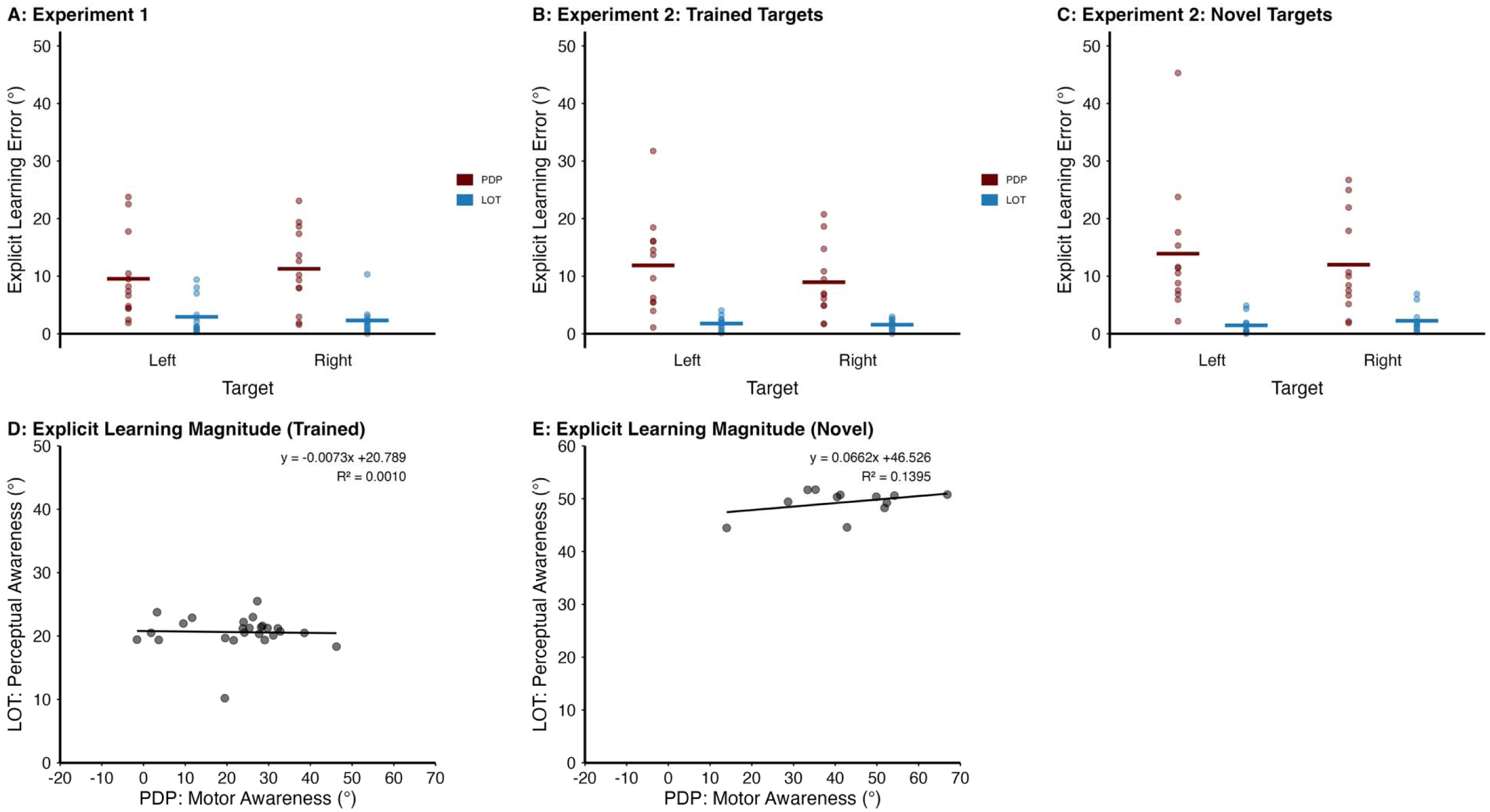
Absolute error in explicit learning across assessment methods and targets (A, B and C) and correlations between PDP and LOT assessments of awareness for the MR group (D and E). A, B and C show group-level (dark, bold line) and individual (circles) absolute error of explicit learning established via the Process Dissociation Procedure (PDP) and the Line Orienting task (LOT). **A:** Absolute error of explicit learning towards the trained left and right targets assessed via the PDP (maroon) and LOT (blue) in Experiment 1 and **B:** Experiment 2. **C:** Absolute error of explicit learning towards the novel left and right targets assessed via the PDP (maroon) and LOT (blue) in Experiment 2. **D:** Correlation between PDP and LOT measures of explicit learning for the trained targets in Experiment 1 and Experiment 2. **E:** Correlation between PDP and LOT measures of explicit learning for the novel targets in Experiment 2.

Figure 5D and Figure 5E depict correlations between motor awareness (PDP) and perceptual awareness (LOT) at the trained and novel targets, respectively. In summary, we found no significant correlations between absolute errors in explicit learning when assessed via the PDP and LOT at the trained targets (y = -0.0073x + 20.789, R² = 0.0010, *p* = 0.879) or at the novel targets (y = 0.0662x + 46.526, R² = 0.1395, *p* = 0.232).

## Discussion

The present study asked whether explicit learning in the form of motor and perceptual awareness are engaged when learning to reach with a small MR distortion. Motor and perceptual awareness after learning to reach with an MR distortion were compared to motor and perceptual awareness after learning to reach with a small visuomotor rotation (VR). Motor awareness was assessed using the process dissociation procedure (PDP), in which participants reached while using any movement strategies they had learned. Perceptual awareness was assessed by asking participants to complete a line orienting task (LOT), in which they indicated their intended aiming direction prior to movement. We additionally tested whether both motor and perceptual awareness generalized to novel targets. Across two Experiments, learning to reach with an MR distortion engaged both motor and perceptual awareness. In contrast, learning to reach with a VR distortion did not show evidence of motor or perceptual awareness, consistent with prior research suggesting that explicit processes have minimal contributions when learning to reach with a small VR distortion (i.e., cursor rotation magnitude < 30°; Modchalingam et al., 2019; Neville & Cressman, 2018; Werner et al., 2015). Further, motor and perceptual awareness engaged following reaches with an MR generalized to novel targets. The magnitude of this awareness differed across assessment methods, suggesting that motor and perceptual assessments index dissociable components of explicit learning rather than a single unitary process. Together, these findings suggest that learning to reach with an MR distortion engages different processes compared to learning to reach with a VR distortion. Learning to reach with an MR distortion is supported by explicit processes, while learning to reach with a VR distortion is supported by implicit (i.e., unconscious) processes.

Participants who learned to reach with the MR distortion consistently expressed both forms of explicit learning, as evidenced by their performance on the PDP and LOT assessment trials, and this explicit learning generalized to novel targets. Motor and perceptual awareness engaged in learning to reach with an MR distortion were dissociable in two ways. First, when awareness was assessed at the trained targets, the magnitudes of awareness assessed via the PDP and LOT were not significantly correlated (Figure 5C), indicating that the degree to which participants expressed motor awareness was largely independent of the degree to which they expressed perceptual awareness. This relationship was also absent at the novel targets in Experiment 2 (Figure 5E), suggesting a lack of relationship between the two types of awareness. Second, the two measures differed in their absolute errors of explicit learning: perceptual awareness, assessed via the LOT, had lower absolute errors than motor awareness, assessed via the PDP. An explanation for this could be that the LOT requires participants to report their intended aiming direction prior to movement execution whereas the PDP requires participants to express awareness of their reaching strategy during movement, incorporating additional sources of variability arising from motor execution and online control (Shadmehr & Mussa-ivaldi, 1994; Wolpert & Kawato, 1998).

Overall, the differences observed between the PDP and LOT when establishing motor and perceptual awareness in learning to reach with an MR distortion align with prior work in VR tasks demonstrating that explicit learning can manifest differently depending on manner of assessment (Maresch et al., 2020, 2021). This dissociation between assessments further suggests that explicit learning involves distinct cognitive strategies (Heirani Moghaddam et al., 2021; Maresch et al., 2020, 2021; McDougle & Taylor, 2019). Consistent with our results, prior studies employing a VR distortion have demonstrated that perceptual measures of explicit strategy tend to account for a larger proportion of behavioural change during learning than do measures derived from motor tasks (McDougle & Taylor, 2019; Taylor et al., 2014).

Further evidence for this dissociation between perceptual and motor awareness is provided by participants who did not learn to reach with the MR distortion (MR-NL group; see Supplementary File). Across Experiments, non-learners were observed only in the MR group and not in the VR group, indicating that failure to learn to reach with the distortion was specific to the mirrored mapping. These participants (MR-NL group) often exhibited motor and/or perceptual awareness, yet the magnitude of these processes was reduced, and the absolute error of explicit learning was greater than that of participants who successfully learned to reach with the MR distortion. The absolute error of explicit learning in the MR-NL group was larger than the MR group across experiments at the trained targets and was especially pronounced when awareness was assessed using the PDP. Thus, failure to learn to reach with the MR distortion was not marked by an absence of explicit processes, but by a reduced magnitude and greater error of explicit learning. These findings suggest that engaging explicit processes with high absolute error, such as implementing an imprecise “move opposite” rule, is insufficient to drive MR learning and that explicit processes must be expressed with sufficiently low absolute error to support learning to reach with an MR distortion.

A different pattern emerged when participants learned to reach with the VR distortion. Consistent with previous work, learning to reach with a VR distortion relied on implicit processes (Modchalingam et al., 2019; Neville & Cressman, 2018; Werner et al., 2015). We found that implicit processes were engaged following learning to reach with the VR distortion, using both PDP and LOT assessments. However, implicit processes established in learning to reach with the VR distortion did not generalize to novel targets. This is in line with previous studies demonstrating that implicit learning, when learning to reach with a VR distortion, is typically characterized by a relatively narrow generalization pattern across target directions and contexts (Cressman & Henriques, 2015; Krakauer et al., 2000).

In contrast, learning to reach with an MR distortion showed no reliable evidence of implicit engagement across experiments or PDP and LOT assessments. Although a small effect was observed in Experiment 2 at the trained targets when implicit learning was assessed with the LOT, this effect did not generalize to novel targets, was not replicated across experiments, and was absent when assessed with the PDP. Because the LOT requires participants to first orient a visual line before executing their reach, the observed effect likely reflects a transient bias introduced by the immediately preceding perceptual judgment rather than an engagement of implicit processes.

Across both experiments, MR participants exhibited longer reaction times (RTs) during learning trials relative to VR participants. Previous work has suggested that prolonged RTs are associated with the engagement of explicit, strategic processes and are thought to reflect the time required for additional planning demands that may accompany the engagement of explicit processes (McDougle & Taylor, 2019; Schween et al., 2020; Taylor & Ivry, 2011). In contrast, learning to reach with a VR distortion is typically characterized by shorter RTs, consistent with a greater reliance on implicit learning. Telgen et al. (2014) similarly reported that longer RTs were associated with improved performance during learning to reach with an MR distortion but not during learning to reach with a VR distortion, further supporting the interpretation that RT reflects differential planning demands across MR and VR environments. In the present study, the elevated RTs observed for the MR group are interpreted as a behavioural signature of increased cognitive involvement rather than as a direct measure of explicit processes. Unlike learning to reach with a VR distortion, in which the cursor distortion involves a consistent rotation of movement direction, learning to reach with an MR distortion requires the formation and application of target-specific rule-based mappings (e.g., implementing a “move opposite” transformation that differs across target locations). This added computational burden likely requires additional planning time on each trial. Thus, the RT costs observed for the MR participants are consistent with the engagement of explicit processes indexed by the PDP and LOT measures.

Beyond the engagement of added planning costs, explicit learning has also been shown to generalize to new contexts (Bouchard & Cressman, 2021; Heuer & Hegele, 2011; Werner et al., 2019). We found that motor and perceptual awareness generalized to novel targets in our MR group. MR learners maintained comparable absolute errors of explicit learning when reaching to trained and novel targets, suggesting consistent application of the learned mapping beyond the trained targets. Prior work using large VR distortions has similarly demonstrated that when explicit learning dominates, generalization is broader than when learning is driven primarily by implicit processes (Bouchard & Cressman, 2021; Heuer & Hegele, 2011; Werner et al., 2019). For example, studies that isolate explicit processes during learning to reach with a large VR distortion have shown generalization of explicit processes across the workspace (Bond & Taylor, 2015) and the opposite limb (Bouchard & Cressman, 2021; Heuer & Hegele, 2011). Although learning to reach with large VR distortions also engages implicit processes, the implicit component exhibits relatively narrow generalization (Cressman & Henriques, 2015; Krakauer et al., 2000). In contrast, learning to reach with a small MR distortion in the present study showed no reliable evidence of implicit engagement and was instead characterized by explicit processes that generalized to novel targets. This pattern is consistent with the acquisition of a structural rule for the mirrored mapping (i.e., a “move opposite” transformation) that can be flexibly applied when new target locations are introduced.

## Conclusion

Our results indicate that learning to reach with a small MR distortion engages dissociable explicit processes. Although both perceptual and motor awareness were present in MR learners, their presence alone was not sufficient to drive learning to reach with an MR distortion, as evident in the MR-NL participants. Instead, successful MR learning occurred only when explicit processes were engaged with low absolute error. In addition, learning to reach with an MR distortion contrasted with learning to reach with a VR distortion, which was characterized by implicit engagement and did not generalize to novel targets. In line with this distinction, generalization of learning was observed only for participants in the MR group. Together, these findings indicate that reaching with MR versus VR distortions recruit fundamentally different learning processes and yield distinct patterns of generalization.

## Acknowledgements

This research was supported by a Discovery Grant from the Natural Sciences and Engineering Research Council of Canada (NSERC) awarded to Gerome A. Manson and Erin K. Cressman. We thank the participants who volunteered to take part in this study. We also thank members of the Sensorimotor Exploration Lab for their assistance with data collection and helpful discussions throughout the project.

## Author contributions (CRediT)

Conceptualization: Sarvenaz Heirani Moghaddam, Erin K. Cressman, Gerome A. Manson

Data curation: Sarvenaz Heirani Moghaddam

Formal analysis: Sarvenaz Heirani Moghaddam

Funding acquisition: Erin K. Cressman

Investigation: Sarvenaz Heirani Moghaddam

Methodology: Sarvenaz Heirani Moghaddam, Erin K. Cressman, Gerome A. Manson

Project administration: Sarvenaz Heirani Moghaddam

Resources: Erin K. Cressman, Gerome A. Manson

Software: Gerome A. Manson

Supervision: Erin K. Cressman, Gerome A. Manson

Validation: Erin K. Cressman, Gerome A. Manson

Visualization: Sarvenaz Heirani Moghaddam

Writing – original draft: Sarvenaz Heirani Moghaddam

Writing – review & editing: Sarvenaz Heirani Moghaddam, Erin K. Cressman, Gerome A. Manson

