## Supplementary file for "Engagement of motor and perceptual awareness when learning to reach with mirror reversed feedback"

### 1.1. **Supplementary File**

#### 1.1.1. **Experiment 2 Learning**

The results below are from Experiment 2, in which 40 participants (29 females) were assigned to the MR group (n = 20) or the VR group (n = 20), with VR-CW and VR-CCW subgroups consisting of 10 participants each. Only participants who successfully learned to reach with the MR distortion were included in this file (12 MR participants in Experiment 2). Overall, the pattern of results was similar to that of Experiment 1, reported in the main manuscript.

With respect to learning, ANOVA revealed a significant main effect of time (F(1, 29) = 25.819, p < 0.001, *η_p_²* = 0.471), and a significant group × time interaction (F(2, 29) = 5.111, p = 0.013, *η_p_²* = 0.261). Post hoc analysis indicated that, for the MR group, early AEs toward the right and left targets were similar (p = 1.000). Further, for the MR group, early AEs toward the left target were significantly smaller than late AEs toward the same target (p < 0.001). Additionally, early AEs toward the left target for the MR group were significantly smaller than both early and late AEs in the VR-CW group and the VR-CCW group (all *p* < 0.05). No early–late differences in AE were observed for the VR-CW or VR-CCW groups (all *p* > 0.05). Importantly, late AEs in the learning block did not differ across groups (all p > 0.05; Figure S1A & B).

AE variability across groups in Experiment 2 revealed a significant main effect of group (F(2, 29) = 49.767, p < 0.001, *η_p_²* = 0.774), and a significant interaction between target position × time (F(1, 29) = 0.387, p = 0.002, *η_p_²* = 0.291). Post hoc analysis revealed that AE variability was significantly greater for the MR group than both the VR-CW group and the VR-CCW group (all p < 0.001). No significant difference in AE variability was observed between the VR-CW and VR-CCW groups (p = 1.000; see Figure S1B). Despite the significant interaction, post hoc analyses did not reveal differences related to target position or time (all p > 0.05). AE variability remained elevated for the MR group in both early and late trials in the learning block compared to the VR groups (Figure S1C).

For RT, we found a significant main effect of group (F(2, 29) = 22.021, p < 0.001, *η_p_²* = 0.603), with post hoc analyses indicating that the MR group took significantly longer to initiate their reaches compared to participants in the VR-CCW group and the VR-CW group (all p < 0.001). RTs were similar between the VR-CW and VR-CCW groups (p = 1.000; Figure S1D).

MTs during the learning block were analyzed to confirm adherence to the 600 ms movement criterion. ANOVA revealed a significant main effect of group (F(2, 29) = 7.213, p = 0.003, *η_p_²* = 0.332), and a significant interaction between group × target position × time (F(2, 29) = 4.705, p < 0.001, *η_p_²* = 0.245). Overall, MTs were shorter for the MR and VR-CCW groups than the VR-CW group (all p < 0.05; Figure S1E). Post hoc analyses revealed that MTs for the VR-CW group, early and late in the learning block, were significantly longer than MTs for both the VR-CCW group and the MR group (all p < 0.05) at both time points. In addition, late MTs for the MR group towards the left target were shorter than late MTs in the VR-CW group towards the left target (p < 0.05). Despite these, group and target specific differences, all movements were completed well within the 600 ms limit; mean late MTs across groups ranged from 194–264 ms.

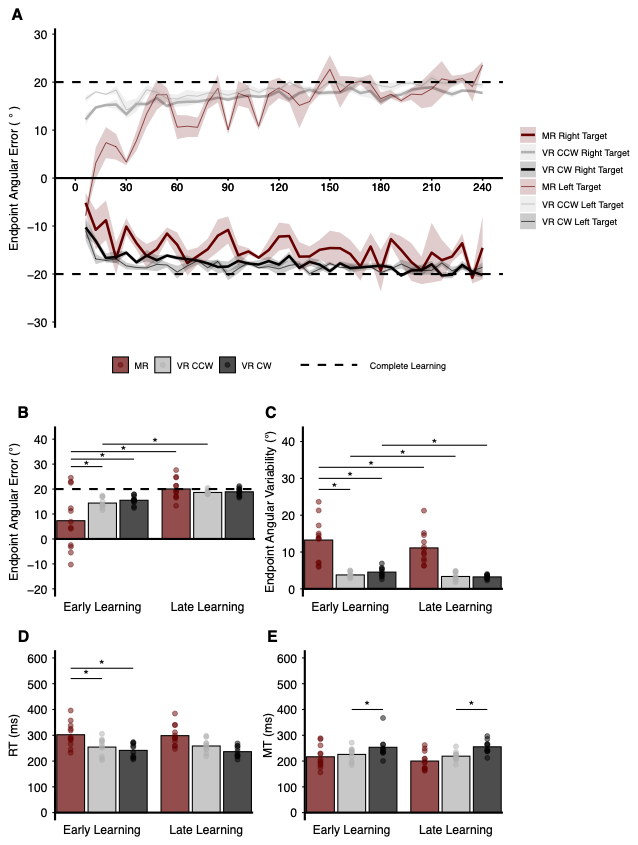

**Figure S1.** Learning performance across groups in Experiment 2. **A:** Learning curves for each group across trials. Each point represents the mean endpoint angular error (AE, in degrees) averaged across three consecutive trials in the learning block. Bold solid lines reflect reaches toward the right target, while thin solid lines indicate reaches toward the left target. Shaded regions represent the standard error of the mean (SEM). Dashed lines indicate the expected reach adjustments for complete learning: y = −20° for the right target, and y = 20° for the left target. Group colors: MR (maroon), VR-CCW (gray), and VR-CW (black). **B:** Absolute AE, **C:** Variability in AE (standard deviation, SD), **D:** Reaction time (RT), and **E:** Movement time (MT) data for each group during early and late learning. Individual participant data are represented as circles. Asterisks denote significant differences between groups at the same time, or within a group across times (*p* ≤ 0.05).

#### 1.1.2. **MR non-learners (MR-NL) subgroup - Learning**

Participants in the MR group were identified as non-learners if their reaching performance during the learning block did not differ from the baseline block. Two criteria were used to classify participants as non-learners: (a) the mean angular error (AE) on late learning trials for either target remained within 3 standard deviations of a participant’s late baseline AE for the same target, and/or (b) reaches were consistently made in the incorrect direction (see Table S1). Applying these criteria, 6 participants in Experiment 1 and 7 participants in Experiment 2 were categorized as non-learners (MR-NL). Follow-up analyses, conducted with the same statistical approach as described in the main manuscript, confirmed that the MR-NL subgroup’s mean AE (normalized to baseline) did not differ between early and late learning (Experiment 1 and Experiment 2: both *p* = 1.000; see Figure S2A & B). These results further verified that these participants did not demonstrate motor learning.

We compared AE variability, movement time (MT), and reaction time (RT) between MR-learners (MR-L) and MR-NL using a 2 group (MR-L, MR-NL) x 2 target position (right, left) x 2 time (early, late) mixed ANOVA, with repeated measures (RM) on the last two factors.

**Experiment 1** For AE variability, we observed significant interactions between group x time (*F*(1,17) = 5.229, *p* = 0.035, *η_p_²* = 0.235), and target position x time (*F*(1,17) = 5.081, *p* = 0.038, *η_p_²* = 0.230). Post hoc analyses indicated that AE variability significantly decreased from early to late learning only for the MR-L group, but this change was not observed for the MR-NL group (Figure S2E). For RT we did not observe any significant main effects or interactions, suggesting no differences in RT between MR-L and MR-NL participants (*F*(1,17) = 0.027, *p* = 0.872, *η_p_²* = 0.0009; Figure S2F). For MT, we found a significant main effect of target position (*F*(1,17) = 15.514, *p* = 0.001, *η_p_²* = 0.447), and a significant interaction between target position x time (*F*(1,17) = 11.738, *p* = 0.003, *η_p_²* = 0.408). Post hoc analyses indicated that the early reaches to the right target had a shorter MT compared to early reaches to the left target (Figure S2G). Overall, in Experiment 1, no group differences were observed between MR-L and MR-NL groups except that AEs for the MR-L group were less variable by the end of the learning block, a pattern that was not observed for the MR-NL group.

**Experiment 2** For AE variability, ANOVA revealed significant interactions between target x time (*F*(1,17) = 6.534, *p* = 0.020, *η²* = 0.278), and group x target position x time (*F*(1,17) = 4.922, *p* = 0.040, *η²* = 0.225). Post hoc analyses did not indicate any significant differences between the MR-L and MR-NL groups (Figure S2H). For RT (Figure S2I) and MT (Figure S2J), there were no significant effects or interactions between the MR-L and MR-NL groups (all *p* > 0.05). Overall, in Experiment 2, there were no differences between the MR-L and MR-NL groups with respect to AE variability, RT and MT.

**Table S1.** Summary of MR-NL participants in Experiment 1 and Experiment 2. Values reflect mean angular error (AE; degrees) in the learning block relative to the baseline block. Each row represents an individual participant’s average AE for early and late learning trials to both right and left targets. Negative values indicate movements in the incorrect direction. The final column specifies the rationale for classifying each participant in the MR-NL subgroup. Participants were classified as learners only if they adjusted reaches to both targets in the correct direction.

| Experiment | Participant | Left Target (°) | | Right Target (°) | | Rationale for designation as  MR-NL |
| --- | --- | --- | --- | --- | --- | --- |
|  |  | **Early** | **Late** | **Early** | **Late** |  |
| 1 | MR_NL01 | -6.4 | -36.2 | 0.5 | 1.9 | Right target AEs not different from baseline; Wrong direction for left target |
|  | MR_NL02 | -22.1 | 28.1 | 4.8 | -15.5 | Wrong direction for right target |
|  | MR_NL03 | 16.5 | 30.5 | 13.9 | -6.3 | Wrong direction for right target |
|  | MR_NL04 | -15.8 | 19.5 | 6.5 | 0.4 | Right target AEs not different from baseline |
|  | MR_NL05 | 3.6 | 6.9 | -12.0 | -16.0 | Wrong direction for right target |
|  | MR_NL06 | 17.5 | 25.1 | -10.2 | -0.1 | Right target AEs not different from baseline |
| 2 | MR_NL01 | -23.0 | 10.2 | 25.0 | -13.2 | Wrong direction for right target |
|  | MR_NL02 | 21.1 | -35.7 | -24.8 | -10.0 | Wrong direction for both targets |
|  | MR_NL03 | 3.2 | 22.1 | 17.8 | -7.9 | Wrong direction for right target |
|  | MR_NL04 | 0.0 | 34.7 | 0.5 | -4.0 | Wrong direction for right target |
|  | MR_NL05 | 9.5 | 23.7 | 6.7 | -4.3 | Wrong direction for right target |
|  | MR_NL06 | -0.5 | -12.2 | 0.3 | 18.3 | Wrong direction for left target |
|  | MR_NL07 | 10.4 | 10.4 | -1.8 | 1.2 | Right target AEs not different from baseline |

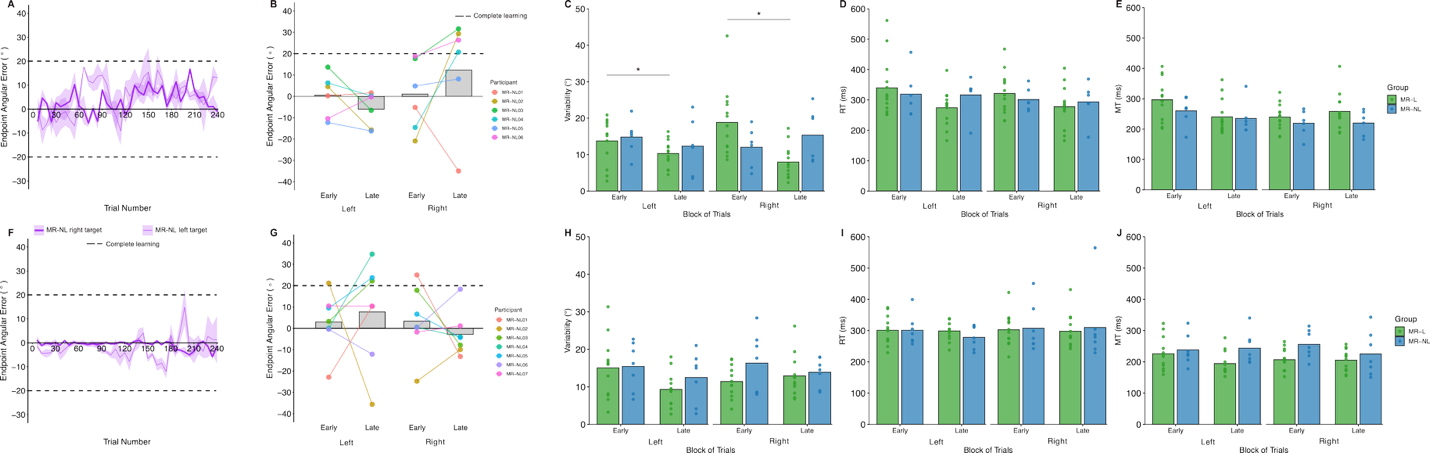

**Figure S2.** MR-NL participants’ learning data in Experiment 1 and Experiment 2. **A:** Learning curve for MR-NL participants in Experiment 1 and **B:** Individual data for the MR-NL group early and late in the learning block in Experiment 1. **C-E:** AE Variability (SD), reaction time (RT) and movement time (MT) for the MR-L (green) and MR-NL (blue) groups in Experiment 1. Bottom row (**F-J)** the same data is presented for Experiment 2. Individual participant data are represented as circles. Asterisks denote significant differences between groups at the same time, or within a group across times (p ≤ 0.05).

#### 1.1.3. **MR-NL subgroup - Explicit Learning**

We compared the magnitude of explicit learning (E_Motor Awareness (PDP)_ and E_Perceptual Awareness (LOT)_) and the absolute error of explicit learning between participants who learned to reach with the MR distortion (MR-L) and those who did not (MR-NL). For each dependent variable (magnitude of explicit learning and absolute error of explicit learning), three mixed analyses of variance (ANOVAs) with repeated measures (RM) on the assessment type were conducted. Specifically, in Experiment 1, magnitude and absolute error were each analyzed in a 2 group (MR-L, MR-NL) × 2 assessment type (PDP, LOT) mixed ANOVA with RM on the last factor. In Experiment 2, separate 2 group × 2 assessment type mixed ANOVAs with RM on the last factor were performed for the trained targets and the novel targets. Thus, for each experiment and target type, the magnitude of explicit learning and its absolute error were evaluated as a function of group and assessment method.

Figure S3A and B display the magnitude of explicit learning for the MR-L participants compared to MR-NL participants. In Experiment 1, a significant main effect of group was observed (*F*(1,17) = 11.333, *p* = 0.004, *η_p_²* = 0.400). Post hoc analyses indicated that the magnitude of explicit learning was larger in the MR-L group compared to the MR-NL group (*p* < 0.05; MR-L: PDP M = 17.0° ± 11.9°, LOT M = 20.3° ± 3.6°; MR-NL: PDP M = 1.7° ± 12.5°, LOT M = 9.6° ± 10.4°). In Experiment 2, at the trained targets, a significant main effect of assessment type (*F*(1,17) = 5.077, *p* = 0.038, *η_p_²* = 0.230) was found, such that motor awareness assessed using the PDP had a larger magnitude than perceptual awareness assessed via the LOT (*p* < 0.05). In Experiment 2, at the novel targets, significant main effects of group (*F*(1,17) = 6.118, *p* = 0.024, *η_p_²* = 0.267) and assessment type (*F*(1,17) = 9.613, *p* = 0.006, *η_p_²* = 0.361) were observed. Post hoc analyses revealed that the MR-L group had larger magnitude of explicit learning compared to the MR-NL group at the novel targets (*p* < 0.05). In addition, perceptual awareness overall had a larger magnitude than motor awareness at the novel targets (*p* < 0.05).

Figure S3C and D illustrate absolute errors in explicit learning for MR-L and MR-NL participants. In Experiment 1, significant main effects of group (*F*(1,17) = 5.746, *p* = 0.028, *η_p_²* = 0.253) and assessment type (*F*(1,17) = 23.109, *p* < 0.001, *η_p_²* = 0.576) were observed, with no significant group × assessment type interaction (*F*(1,17) = 5.746, *p* = 0.028, *η_p_²* = 0.253). Post hoc analyses indicated larger absolute errors of explicit learning for the MR-NL group than for the MR-L group (*p* < 0.05), and larger absolute errors when assessed using the PDP compared to the LOT (*p* < 0.05). Similarly, in Experiment 2 at the trained targets, significant main effects of group (*F*(1,17) = 4.671, *p* = 0.045, *η_p_²* = 0.216) and assessment type (*F*(1,17) = 40.978, *p* < 0.001, *η_p_²* = 0.707) were found. At the novel targets, significant main effects of group (*F*(1,17) = 8.841, *p* = 0.009, *η_p_²* = 0.342) and assessment type (*F*(1,17) = 45.250, *p* < 0.001, *η_p_²* = 0.727) were also observed. Consistent with Experiment 1, post hoc analyses in Experiment 2 for both trained and novel targets revealed larger absolute errors of explicit learning for the MR-NL group compared to the MR-L group (*p* < 0.05) and larger absolute errors when assessed using the PDP compared to the LOT (*p* < 0.05).

Together, these results indicate that although MR-NL participants engaged explicit processes, those processes were expressed at a lower magnitude and with greater absolute error than in MR-L participants.

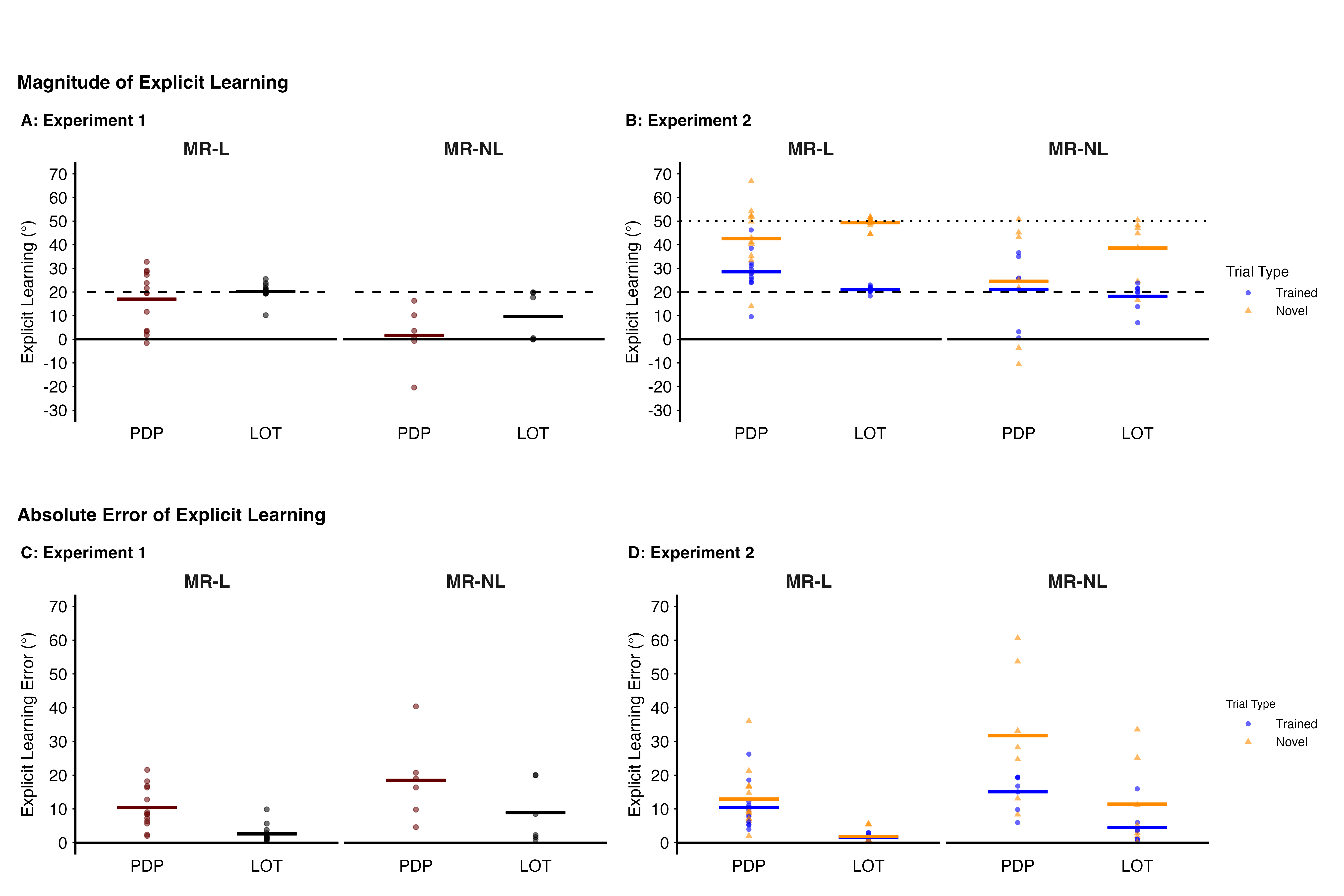

**Figure S3.** Explicit learning magnitude and absolute errors when assessed via the PDP and LOT in MR learners (MR-L) and MR non-learners (MR-NL). All figures show group-level (dark, bold line) and individual explicit learning (circles) and absolute error of explicit learning (circles) established via the Process Dissociation Procedure (PDP) and the Line Orienting task (LOT). **A and B:** Explicit learning magnitude in Experiment 1 (A) and Experiment 2 (B) for the trained targets, when assessed via the PDP and LOT. Dashed horizontal lines indicate the learned criterion for the trained (thick dashed lines) and novel (thin dashed line) targets. In Experiment 2, data are shown separately for trained (blue circles) and novel (orange triangles) targets. **C and D:** Absolute errors in explicit learning in Experiment 1 (C) and Experiment 2 (D) assessed via the PDP and LOT. Absolute errors are plotted separately for MR-L and MR-NL groups, and for trained and novel targets in Experiment 2.
